## Supplementary_files for "Optimized Mass Spectrometry Detection of Thyroid Hormones and Polar Metabolites in Rodent Cerebrospinal Fluid"

Figure S1. Limit of detection and linearity for the individual thyroxin and triiodothyronine standards for the indicated chromatographic methods: "ZIC-pHILIC": ZIC-pHILIC 150 × 2.1 mm (5 µm particle size, EMD Millipore; "Accucore Amide": Accucore™ 150 Amide HILIC (150x3 mm, 2.6 mm particle size; Thermo Fisher Scientific) ; "LUNA NH2": Luna® 3 µm NH2 100 Å, LC Column (150x2 mm, 3 µm particle size; Phenomenex, 00F-4377-B0). Presented is one experiment from two representative dilutions; R2 – goodness of fit; Sy.x - standard deviation of the residuals. "LOD" – limit of detection.

Figure S2 and Figure S3. Spectral information on T3 (S2) and T4 (S3). Four panels representing spectra collected with a consecutive increase in HCD energy: 20, 40, 60, and 80 NCE.

Figure S4. Optimization of reconstitution conditions for T4 and T3 from cerebrospinal fluid (CSF). (a) Rat serum was extracted with a two-phase extraction. Both lipid (bottom) and polar (top) phases from a two-phase extraction were compared, where each was reconstituted in either water or 70% ACN. Normalized peak integration areas are shown as the average and standard deviation for two independent extractions.

Figure S5. Optimization of solvents for extraction of T3 and T4 from adult and embryonic CSF. Single-phase ("MB": methanol based, see methods for further details) vs two-phase (lipid and polar phase respectively) extractions are compared. Normalized peak integration areas are shown as the average and standard deviation for a triplicate adult or embryo CSF collection.

Figure S6. Levels of T3 and T4 between adult and embryo. (a) Comparison of storage conditions and signal stability upon freezing and storage for T3 or T4 levels extracted from CSF from adult mice or embryo as well as adult serum. This data is related to Figure 4A which only shows the "fresh" set of samples. Normalized peak integration areas are shown as the average and standard deviation for at least five CSF collections and four paired plasma collections. (b) Comparison of storage conditions and signal stability upon freezing and storage for mouse CSF and serum T4. Freshly extracted CSF extract was compared to flash-frozen vs flash-frozen and 24h stored. (c-f) Comparison between female and male or embryo fresh and frozen CSF polar metabolites. (c and d) Heatmap analysis showing top 25 changed metabolites from indicated CSF conditions and analysed using HILIC chromatography LC-MS. Detected metabolites were Pareto scaled and log-transformed within the MetaboAnalyst online platform. (e and f) Corresponding volcano plots for the data in (c) and (d) respectively.

**Figure S1**

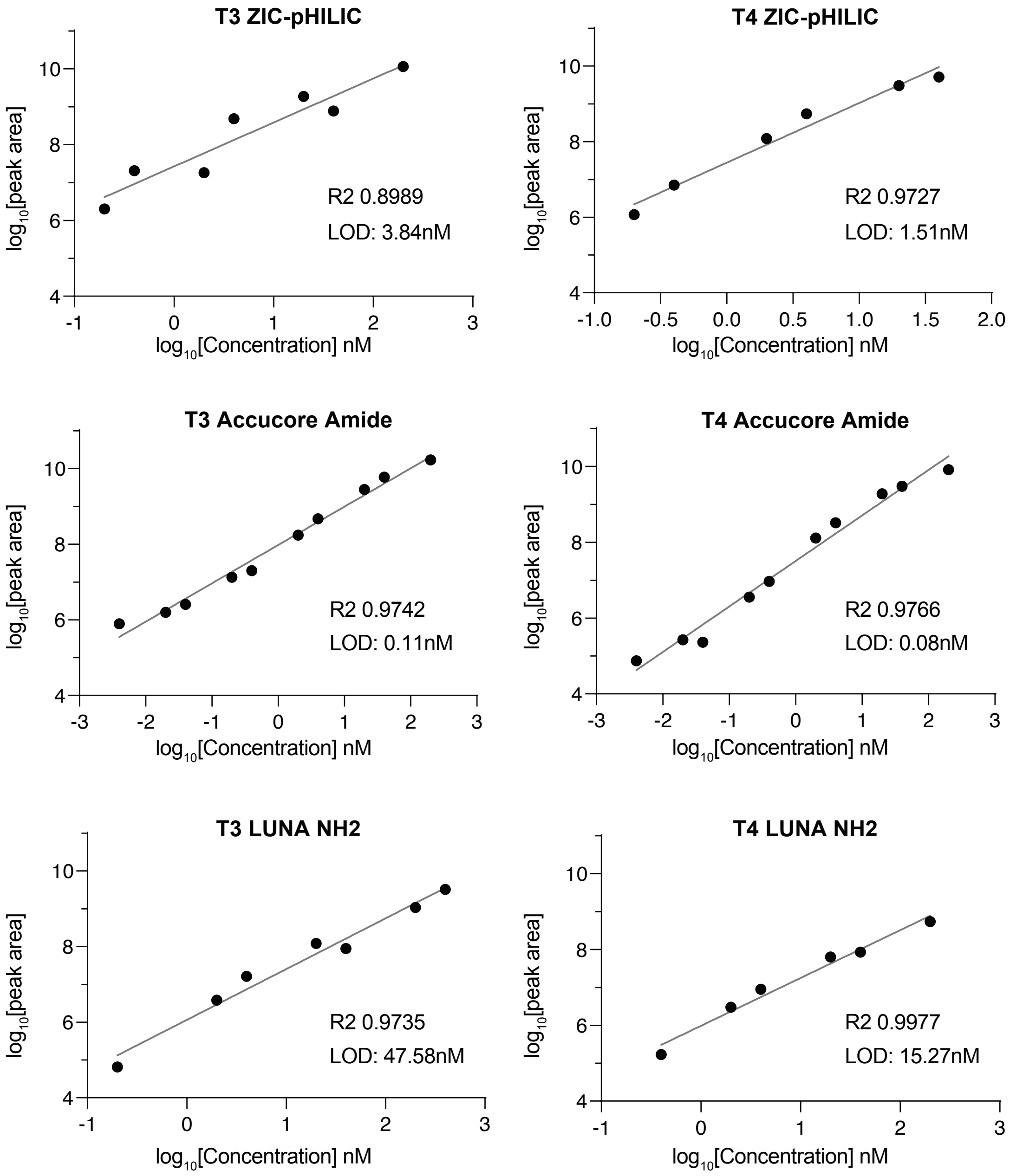

T3 ZIC-pHILIC
R2 0.8989
LOD: 3.84nM
T4 ZIC-pHILIC
R2 0.9727
LOD: 1.51nM
T3 Accucore Amide
R2 0.9742
LOD: 0.11nM
T4 Accucore Amide
R2 0.9766
LOD: 0.08nM
T3 LUNA NH2
R2 0.9735
LOD: 47.58nM
T4 LUNA NH2
R2 0.9977
LOD: 15.27nM
log10[peak area]
log10[Concentration] nM

#### Figure S2

### A

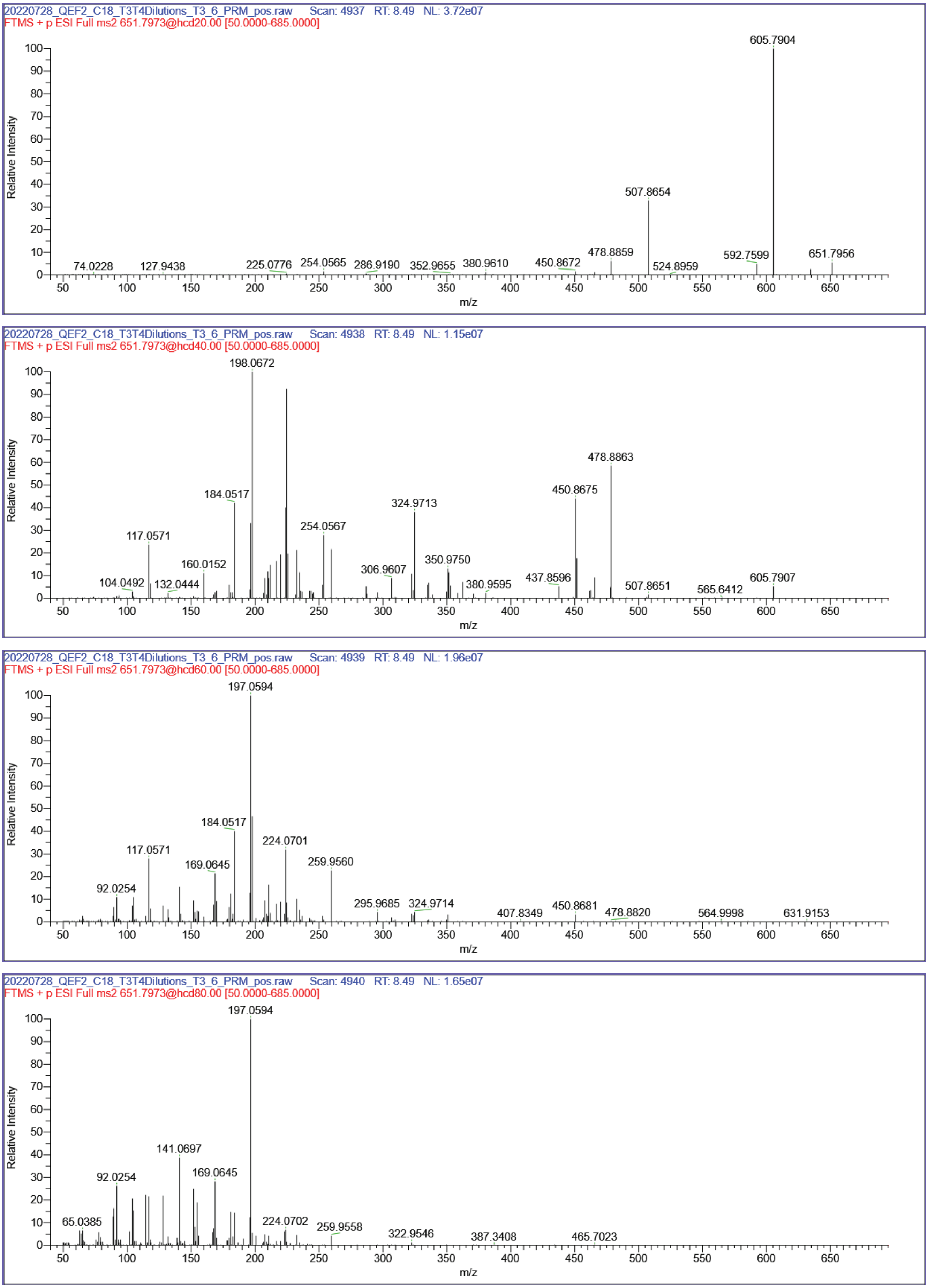

20220728_QEF2_C18_T3T4Dilutions_T3_6_PRM_pos.raw Scan: 4937 RT: 8.49 NL: 3.72e07
FTMS + p ESI Full ms2 651.7973@hcd20.00 [50.0000-685.0000]
605.7904
507.8654
74.0228
127.9438
225.0776
254.0565
286.9190
352.9655
380.9610
450.8672
478.8859
524.8959
592.7599
651.7956
Relative Intensity
m/z
20220728_QEF2_C18_T3T4Dilutions_T3_6_PRM_pos.raw Scan: 4938 RT: 8.49 NL: 1.15e07
FTMS + p ESI Full ms2 651.7973@hcd40.00 [50.0000-685.0000]
198.0672
478.8863
450.8675
184.0517
324.9713
254.0567
117.0571
160.0152
350.9750
306.9607
104.0492
132.0444
380.9595
437.8596
507.8651
565.6412
605.7907
Relative Intensity
m/z
20220728_QEF2_C18_T3T4Dilutions_T3_6_PRM_pos.raw Scan: 4939 RT: 8.49 NL: 1.96e07
FTMS + p ESI Full ms2 651.7973@hcd60.00 [50.0000-685.0000]
197.0594
184.0517
224.0701
117.0571
169.0645
259.9560
92.0254
295.9685
324.9714
407.8349
450.8681
478.8820
564.9998
631.9153
Relative Intensity
m/z
20220728_QEF2_C18_T3T4Dilutions_T3_6_PRM_pos.raw Scan: 4940 RT: 8.49 NL: 1.65e07
FTMS + p ESI Full ms2 651.7973@hcd80.00 [50.0000-685.0000]
197.0594
141.0697
92.0254
169.0645
65.0385
224.0702
259.9558
322.9546
387.3408
465.7023
Relative Intensity
m/z

**Figure S3**

**A**

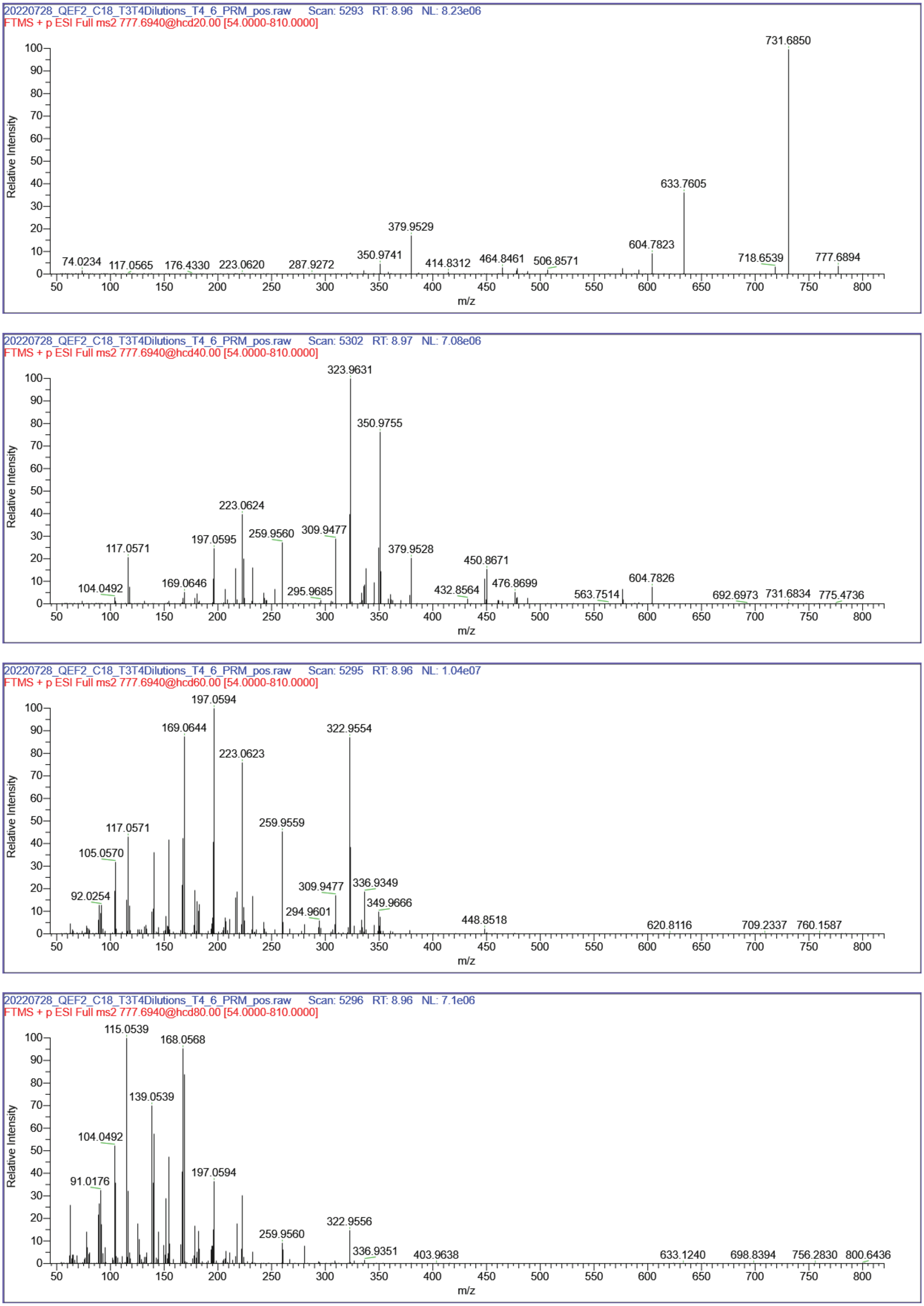

20220728_QEF2_C18_T3T4Dilutions_T4_6_PRM_pos.raw Scan: 5293 RT: 8.96 NL: 8.23e06
FTMS + p ESI Full ms2 777.6940@hcd20.00 [54.0000-810.0000]
731.6850
633.7605
379.9529
604.7823
350.9741
74.0234
117.0565
176.4330
223.0620
287.9272
414.8312
464.8461
506.8571
718.6539
777.6894
Relative Intensity
m/z
20220728_QEF2_C18_T3T4Dilutions_T4_6_PRM_pos.raw Scan: 5302 RT: 8.97 NL: 7.08e06
FTMS + p ESI Full ms2 777.6940@hcd40.00 [54.0000-810.0000]
323.9631
350.9755
223.0624
259.9560
309.9477
197.0595
117.0571
379.9528
450.8671
169.0646
104.0492
295.9685
432.8564
476.8699
604.7826
563.7514
692.6973
731.6834
775.4736
Relative Intensity
m/z
20220728_QEF2_C18_T3T4Dilutions_T4_6_PRM_pos.raw Scan: 5295 RT: 8.96 NL: 1.04e07
FTMS + p ESI Full ms2 777.6940@hcd60.00 [54.0000-810.0000]
197.0594
169.0644
322.9554
223.0623
259.9559
117.0571
105.0570
92.0254
309.9477
336.9349
349.9666
294.9601
448.8518
620.8116
709.2337
760.1587
Relative Intensity
m/z
20220728_QEF2_C18_T3T4Dilutions_T4_6_PRM_pos.raw Scan: 5296 RT: 8.96 NL: 7.1e06
FTMS + p ESI Full ms2 777.6940@hcd80.00 [54.0000-810.0000]
115.0539
168.0568
139.0539
104.0492
91.0176
197.0594
322.9556
259.9560
336.9351
403.9638
633.1240
698.8394
756.2830
800.6436
Relative Intensity
m/z

### Figure S4

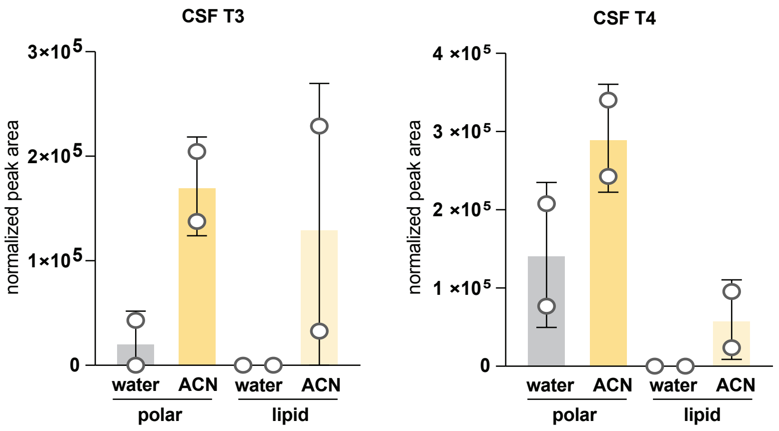
CSF T3
normalized peak area
3×10^5
2×10^5
1×10^5
0
water
ACN
water
ACN
polar
lipid
CSF T4
normalized peak area
4 ×10^5
3 ×10^5
2 ×10^5
1 ×10^5
0
water
ACN
water
ACN
polar
lipid

### Figure S5

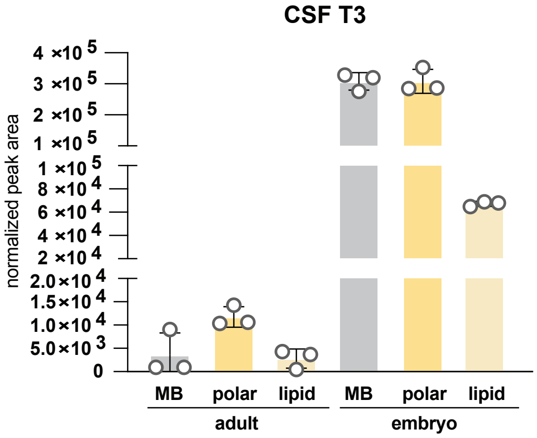
CSF T3
normalized peak area
4 ×10 5
3 ×10 5
2 ×10 5
1 ×10 5
1 ×10 5
8 ×10 4
6 ×10 4
4 ×10 4
2 ×10 4
2.0×10 4
1.5×10 4
1.0×10 4
5.0×10 3
0
MB
polar
lipid
MB
polar
lipid
adult
embryo

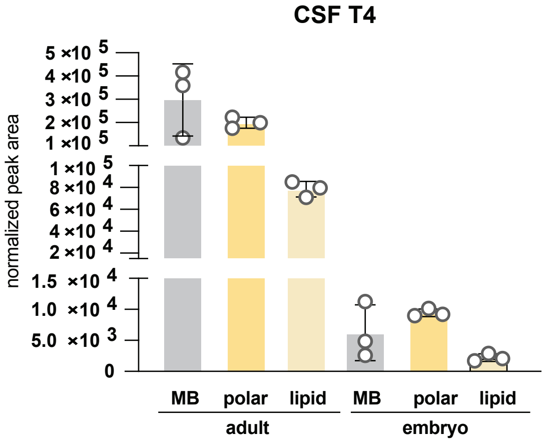
CSF T4
normalized peak area
5 ×10 5
4 ×10 5
3 ×10 5
2 ×10 5
1 ×10 5
1 ×10 5
8 ×10 4
6 ×10 4
4 ×10 4
2 ×10 4
1.5 ×10 4
1.0 ×10 4
5.0 ×10 3
0
MB
polar
lipid
MB
polar
lipid
adult
embryo

Figure S6

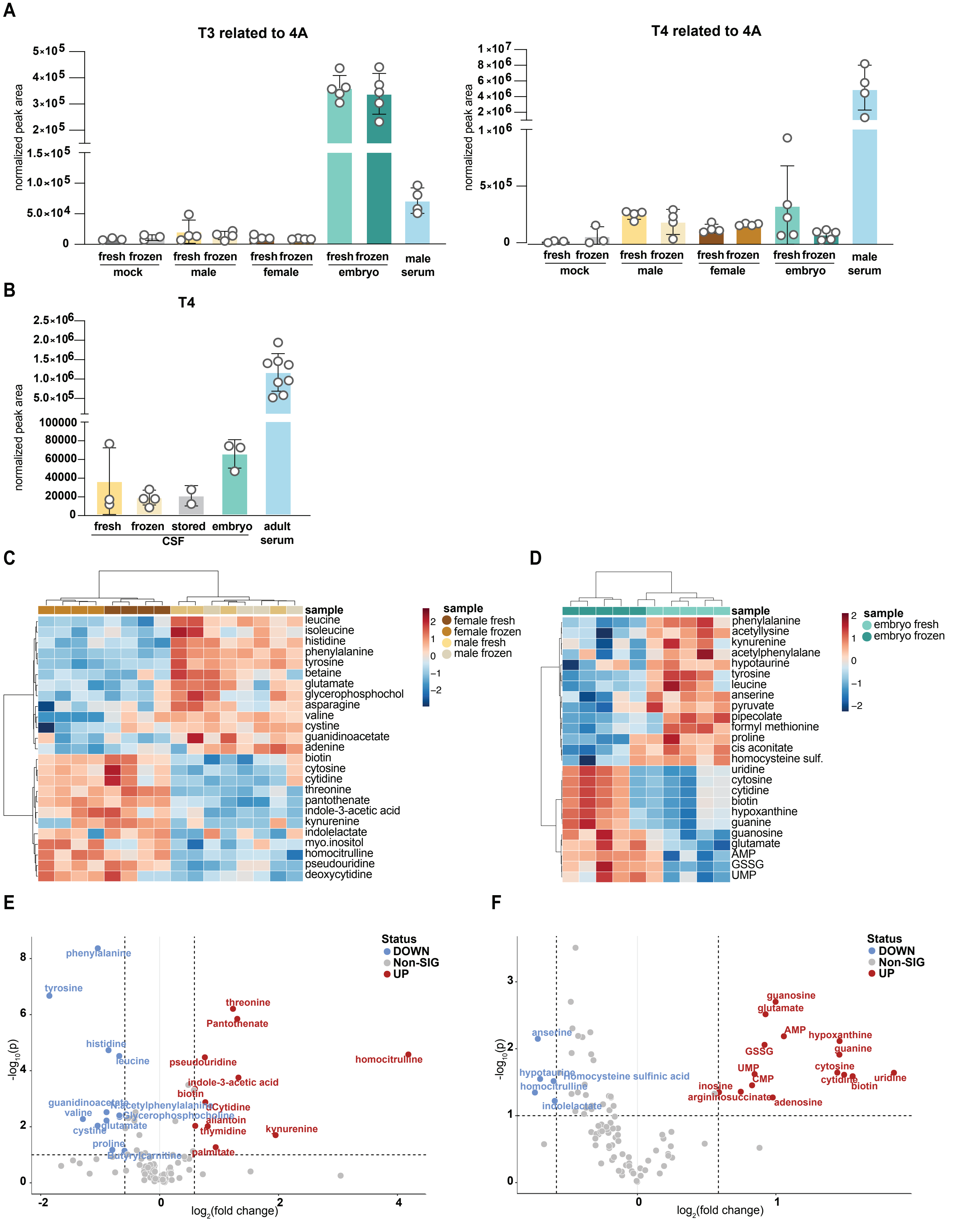
A
T3 related to 4A
normalized peak area
fresh frozen
mock
male
female
embryo
male
serum
T4 related to 4A
B
T4
fresh
frozen
stored
embryo
adult
serum
CSF
C
sample
female fresh
female frozen
male fresh
male frozen
leucine
isoleucine
histidine
phenylalanine
tyrosine
betaine
glutamate
glycerophosphochol
asparagine
valine
cystine
guanidinoacetate
adenine
biotin
cytosine
cytidine
threonine
pantothenate
indole-3-acetic acid
kynurenine
indolelactate
myo.inositol
homocitrulline
pseudouridine
deoxycytidine
D
sample
embryo fresh
embryo frozen
phenylalanine
acetyllysine
kynurenine
acetylphenylalane
hypotaurine
tyrosine
leucine
anserine
pyruvate
pipecolate
formyl methionine
proline
cis aconitate
homocysteine sulf.
uridine
cytosine
cytidine
biotin
hypoxanthine
guanine
guanosine
glutamate
AMP
GSSG
UMP
E
Status
DOWN
Non-SIG
UP
log2(fold change)
-log10(p)
phenylalanine
tyrosine
histidine
leucine
guanidinoacetate
valine
cystine
glutamate
proline
Butyrylcarnitine
N.acetylphenylalanine
Glycerophosphocholine
threonine
Pantothenate
homocitrulline
pseudouridine
indole-3-acetic acid
biotin
dCytidine
allantoin
thymidine
kynurenine
palmitate
F
Status
DOWN
Non-SIG
UP
anserine
hypotaurine
homocitrulline
Homocysteine sulfinic acid
indolelactate
guanosine
glutamate
AMP
hypoxanthine
guanine
GSSG
UMP
CMP
cytosine
cytidine
uridine
biotin
inosine
argininosuccinate
adenosine
